# RNA-seq: primary cells, cell lines and heat stress

**DOI:** 10.1101/013979

**Authors:** Carl J. Schmidt, Elizabeth M. Pritchett, Liang Sun, Richard V.N. Davis, Allen Hubbard, Kalmia E. Kniel, Sarah M. Markland, Qing Wang, Chris Ashwell, Michael Persia, Max F. Rothschild, Susan J. Lamont

**Affiliations:** Department of Animal and Food Sciences, University of Delaware, Newark DE. 19716 Elizabeth M. Pritchett Liang Sun Richard V.N. Davis Allen Hubbard Kalmia E. Kniel Sarah M. Markland Qing Wang; Department of Poultry Science, 147 Scott Hall, NCSU, Raleigh, NC 27696 Chris Ashwell; Department of Animal and Poultry Science, 3060 Litton-Reaves Hall, Virginia Tech, Blacksburg, VA. 24061. Michael Persia; Department of Animal Science, Iowa State University, 2255 Kildee Hall, Ames IA 50011. Max F. Rothschild; Susan J. Lamont

## Abstract

Transcriptome analysis by RNA-seq has emerged as a high-throughput, cost-effective means to evaluate the expression pattern of genes in organisms. Unlike other methods, such as microarrays or quantitative PCR, RNA-seq is a target free method that permits analysis of essentially any RNA that can be amplified from a cell or tissue. At its most basic, RNA-seq can determine individual gene expression levels by counting the number of times a particular transcript was found in the sequence data. Transcript levels can be compared across multiple samples to identify differentially expressed genes and infer differences in biological states between the samples. We have used this approach to examine gene expression patterns in chicken and human cells, with particular interest in determining response to heat stress.

---

Three separate *in vitro* cell culture experiments were conducted using the white leghorn chicken hepatocarcinoma LMH cell line, the primary chicken liver (e15) line and the human colorectal cancer epithelial Caco-2 cell line. Cells were freshly plated at subconfluent density and grown for 18 hours at 37°C. Six biological replicates were prepared for the LMH cells while three biological replicates were prepared for the primary chicken hepatocytes and Caco-2 cells. After 18 hours, half of the plates were transferred to 43°C for 2.5 hours after which RNA was isolated, Illumina RNA-seq libraries prepared (Coble et al. 2014; Sun et al. 2015) and sequenced to a depth of 10 million or more reads by the University of Delaware DNA sequencing facility. In addition, we prepared RNA-seq libraries were prepared from six freshly isolated 7-day old chicken livers.

Sequence data were analyzed using the Tuxedo software suite (Gal4 reference build and annotation) and differentially enriched genes identified as indicated below. Signaling and metabolic pathways affected by enriched genes were identified using Gallus Reactome^**+**^ (http://gallus.reactome.org/) and eGIFT (Tudor et al. 2010).

RNA-seq data were combined using chicken:human orthologs to merge the data (Burt et al. 2009) yielding a total of 5333 pairs. Initially, hierarchical clustering was used to compare overall gene expression patterns across the cells and tissues (Fig.1). Although prepared at two different developmental time points (e15 vs. d7), the gene expression patterns of the primary liver culture and freshly prepared liver samples cluster together. The chicken LMH and human Caco-2 cells segregate into a second cluster. Overall, the clustering indicates that the primary cultured liver cells are more similar to fresh liver samples than the established cell lines. The clustering of the cell lines likely reflects the transformed nature and long-term culturing of the LMH and Caco-2 lines. Comparing iTerms (Tudor et al. 2010) for genes enriched in either the cell line samples or the liver samples (Fig.2) indicates enrichment for genes involved in cell proliferation in the cell lines. In contrast, the primary liver and cultured liver samples are enriched for genes involved in functions associated with primary hepatic metabolic processes such as coagulation, cholesterol metabolism, and bile production. The latter observation suggests also that the short-term *in vitro* cultured liver cells may model responses of the intact liver.

**Figure 1.**
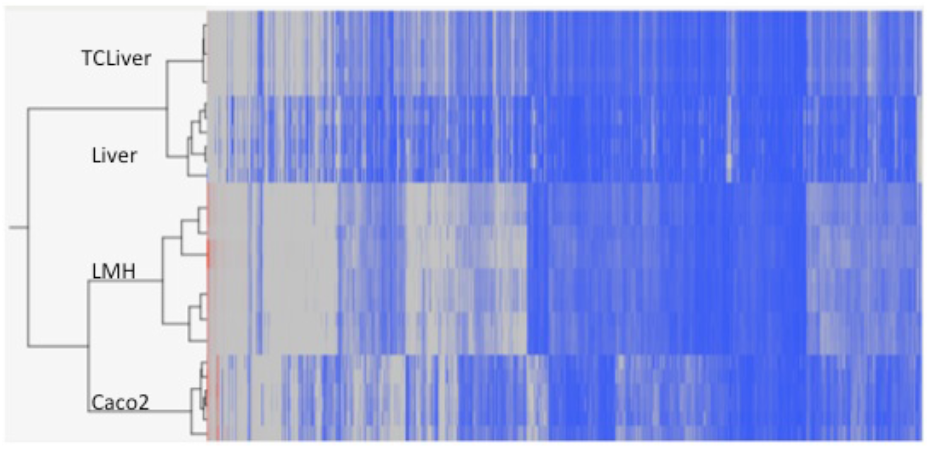
**Hierarchical clustering** of RNA-seq data from primary cultured liver cells (TCLiver), fresh Liver, Chicken white-leghorn hepatocarcioma cell line (LMH) and human colorectal cancer cell line (Caco-2).

**Figure 2.**
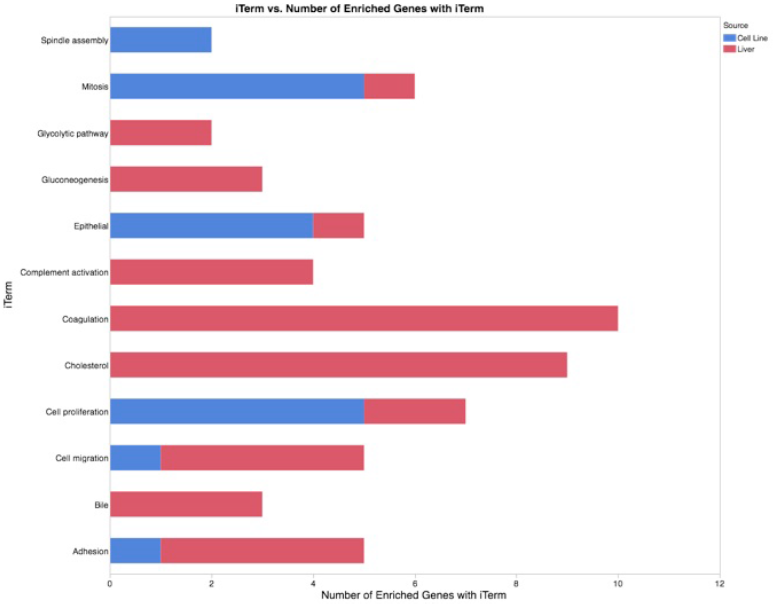
**Stacked Histogram of iTerms** describing biological processes affected by genes enriched in the primary cultured liver samples plus fresh liver (red) compared with genes enriched in the LMH plus Caco-2 cell lines (blue).

## Heat Stress

One of the objectives of our work is identifying pathways responsive to heat stress, with a particular interest in identifying evolutionarily conserved responses. To this end, genes were identified that were responsive to heat stress in the cultured liver, LMH and Caco-2 cells. The genes were then mapped to Gallus Reactome^**+**^ (Sun, manuscript in preparation) to identify signaling and metabolic pathways that were modulated (p value <0.05) by heat stress in all three cell types. A total of 280 genes mapped to Gallus Reactome^**+**^ and affected 22 pathways (Table 1). These pathways impact many cellular activities, including the immune system, basic metabolism, response to growth factors and other stimuli, membrane transport of small molecules along with the integrity of the extracellular matrix. Presumably, these represent a core set of pathways that are fundamental to the overall heat stress response.

**Table 1.** Pathways affected by heat stress in *in vitro* cultured primary cultured liver, LMH and Caco-2 cells.

|  |
| --- |
| **Gallus Reactome plus Pathways** |
| Adaptive Immune System |
| Biological oxidations |
| Collagen formation |
| Degradation of the extracellular matrix |
| Generic Transcription Pathway |
| Innate Immune System |
| Ion channel transport |
| Metabolism of amino acids and derivatives |
| Metabolism of carbohydrates |
| Metabolism of lipids and lipoproteins |
| Peptide hormone metabolism |
| Platelet activation, signaling and aggregation |
| Post-translational protein modification |
| Regulation of Insulin-like Growth Factor (IGF) Transport and Uptake by Insulin-like Growth Factor Binding Proteins (IGFBPs) |
| Regulation of mRNA Stability by Proteins that Bind AU-rich Elements |
| Signaling by ERBB2 |
| Signaling by ERBB4 |
| Signaling by GPCR |
| Signaling by NOTCH |
| Signaling by NGF |
| SLC-mediated transmembrane transport |
| Transmission across Chemical Synapses |

To identify a conserved set of genes that were responsive to heat stress in all three cultured cell types, pairwise comparisons were conducted of the three experiments to identify genes that were regulated in the same direction (up or down) whose difference was statistically significant (p value < 0.05) (Table 2). The transcription levels of 17 genes were differentially regulated by heat stress across all three cell types; 16 were up-regulated by heat stress while one was down-regulated. Eleven of the up-regulated genes function directly as molecular chaperones or as co-chaperones (HSPB8, SERPINH1, BAG3, HSPH1, HSP90AA1, HSPA5, DNAJA1, CHORDC1, HSPA4, HSPA4L and possibly, TPPP). Several of the genes products, such as HSP90AA1, HSPA4, and HSPB8 serve as chaperones for multiple client proteins with broad effects on cellular pathways. Some of these gene products function in specific responses. For example HSPA5 functions during endoplasmic reticulum stress, while SERPINH1 affects collagen folding and assembly. CHORDC1 binds the nucleotide-binding domain of HSP90 when the ADP: ATP ratio is high suggesting CHORDC1 may modulate HSP90 activity as a function of energy balance (Gano and Simon 2010).

**Table 2.** Differentially Expressed Genes responsive to heat stress. P-value for difference between heat-stressed and control samples derived from primary cultured liver (TC_Liver), LMH cells (LMH) and Caco-2 cells followed by the mean values for the control and heat stress RPMK values.

| Symbol | Description | TC_Liver<br>P-value | Caco-2 P-value | LMH P-value | Mean<br>Control<br>TC_Liver | Mean Heat<br>TC_Liver | Mean<br>Control<br>Caco-2 | Mean<br>Heat<br>Caco-2 | Mean<br>Control<br>LMH | Mean<br>Heat<br>LMH |
| --- | --- | --- | --- | --- | --- | --- | --- | --- | --- | --- |
| HSPA5 | Heat Shock 70kDa Protein 5 | 0.0002 | 0.0088 | <.0001 | 418 | 2411 | 265 | 562 | 316 | 1710 |
| HSPB8 | Heat Shock 22kDa Protein 8 | 0.0004 | 0.0145 | 0.0001 | 1.17 | 12.4 | 19.1 | 52.7 | 41.2 | 1484 |
| HSP90AA1 | Heat Shock Protein 90kDa alpha 1 | <.0001 | 0.0423 | <.0001 | 52.4 | 549 | 239 | 943 | 205 | 1369 |
| BAG3 | BCL2-associated athanogene 3 | 0.0002 | 0.0307 | <.0001 | 47.6 | 665 | 14.8 | 198 | 33.8 | 806 |
| HSPH1 | Heat Shock 105kDa/110kDa Protein 1 | <.0001 | 0.0180 | <.0001 | 12.6 | 183 | 43 | 468 | 47.9 | 373 |
| TRA2A | Transformer 2 Alpha | 0.0002 | 0.0135 | <.0001 | 36.5 | 68.9 | 33 | 63.8 | 79.9 | 194 |
| HSPA4L | Heat Shock 70kDa Protein 4 | <.0001 | 0.0417 | 0.0003 | 30.5 | 91.3 | 2.84 | 33.9 | 78.7 | 175 |
| DNAJA1 | DnaJ (Hsp40) | <.0001 | 0.0099 | <.0001 | 28.3 | 114 | 161 | 509 | 54.1 | 171 |
| CHORDC1 | Cysteine and Histidine-rich domain containing 1 | 0.0009 | 0.0366 | <.0001 | 7.76 | 29.6 | 62 | 115 | 39.4 | 124 |
| OTUD1 | OTU Domain Containing 1 | 0.0055 | 0.0146 | 0.0021 | 15.1 | 164 | 8.03 | 21.3 | 14 | 104 |
| HSPA4 | Heat Shock 70kDa Protein 4 | <.0001 | 0.0342 | <.0001 | 18.5 | 66.7 | 65.4 | 117 | 29.1 | 88.8 |
| TPPP | Tubulin Polymerization Promoting Protein | 0.0064 | 0.0451 | 0.0076 | 0.42 | 0.89 | 0.09 | 0.43 | 11.6 | 26.6 |
| SNTB2 | Syntrophin, beta 2 | 0.0018 | 0.0072 | 0.0014 | 2.29 | 4.93 | 4.98 | 9.81 | 6.97 | 15.4 |
| VRK1 | Vaccinia Related Kinase 1 | <.0001 | 0.0106 | 0.0010 | 5.54 | 2.03 | 22.2 | 6.79 | 24.8 | 7.23 |
| SERPINH1 | Serpin Peptidase Inhibitor, clade H -1 | 0.0005 | 0.0346 | <.0001 | 31.1 | 87.8 | 231 | 799 | 0.29 | 2.41 |
| OTOA | Otoancorin | 0.0018 | 0.0289 | <.0001 | 0.14 | 0.96 | 0.02 | 0.12 | 0.53 | 2.19 |
| IMPG2 | Interphotoreceptor Matrix Proteoglycan 2 | 0.0002 | 0.0315 | 0.0024 | 0.11 | 1.09 | 0.06 | 0.12 | 0.39 | 1.14 |

The hypothesis that TPPP may function as a chaperone or co-chaperone arises from several observations. One observation is from this work showing that TPPP transcription is induced in chicken and human cells by heat stress. Furthermore, the TPPP gene product is known to modulate the tubulin network, promoting tubulin polymerization, microtubule acetylation and bundling, along with alpha-synuclein autophagy (Ejlerskov et al. 2013; Tirian et al. 2003; Tokesi et al. 2010). Taken together this evidence suggests that TPPP may function as a chaperone of proteins that interact with the tubulin matrix. While predominately studied in neuronal cells, TPPP may have similar functions in other cells.

Other products encoded by these heat responsive genes may not function as chaperones. OTUD1 is a deubiquitinase that regulates the level of type 1 interferon in human cells and functions as a suppressor of the innate immune response (Kayagaki et al. 2007). The deubiquitinase activity may allow UTUD1 to serve as a broad regulator of the ubiquitin protein degradation pathway during the unfolded protein response of heat stress. TRA2 is a nuclear protein, originally identified as controlling sex specific splicing in Drosophila (Amrein et al. 1990; Baker et al. 1989; Goralski et al. 1989) and controls alternative splicing in vertebrates (Zhang et al. 2007). Possibly, heat stress induction of TRA2 leads to an evolutionarily conserved pattern of alternative splicing (Strasburg and Chiang 2009).

Three of the conserved heat responsive genes function either extracellularly or by interaction with the extracellular matrix. OTOA was originally identified as a protein that links the apical surface of inner ear epithelial cells to the extracellular gel (Zwaenepoel et al. 2002). IMPG2 encodes a hyaluronic binding proteoglycan (Acharya et al. 2000). The final heat responsive up-regulated gene, SNTB2 encodes a member of the syntrophin family that links the cytoskeleton with the extracellular matrix (Ervasti and Campbell 1993). The sole gene down regulated by heat stress across all three cell types was VRK1, Vaccinia related kinase 1. This kinase functions during the cell cycle, and is required for transiting the G1 phase (Valbuena et al. 2008). Down-regulation by heat would likely reduce the rate of cell proliferation, possibly permitting more time to repair stress associated cell damage.

The great depth of reads provided by RNA-seq can challenge assertions that a gene is only expressed in a specific cell or tissue. For example the OTOA gene was originally identified as expressed only in inner ear, while the IMPG2 gene has been identified as specific to retinal and pineal tissue (Acharya et al. 2000). However, we detect low levels of both OTOA and IMPG2 transcripts in all three cell types. Inspection of the RNA-seq coverage for both genes revealed multiple reads mapping to the corresponding transcripts with the preponderance coming from exons (Fig. 3). Also, some of the reads span an intron. Taken together these observations suggest that the detected reads arise from mature transcripts rather than precursor RNAs or genomic contamination. The conclusion is that the reads from these genes in our experiments arose from true OTOA or IMPG2 transcripts. However, there is no current evidence that the OTOA or IMPG2 transcripts give rise to protein. This demonstrates an important current challenge: linking data from high throughput sequencing and proteomics analysis to verify translation of such mRNAs.

**Figure 3.**
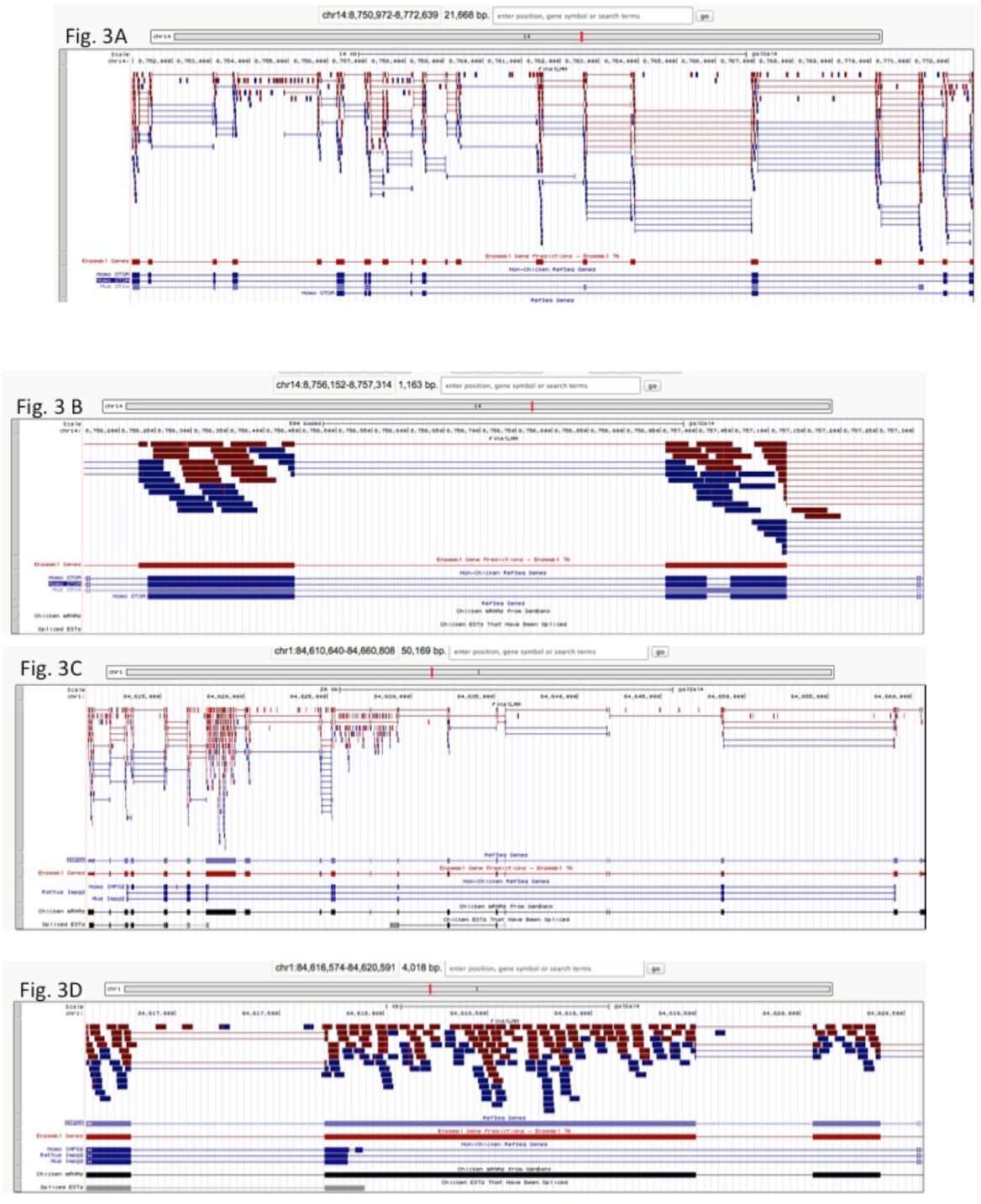
**Track diagrams from UCSC browser** plotting chicken LMH RNA-seq reads from the OTOA (A & B) or IMPG2 (C & d). A and C correspond to the entire gene region while B and D have been zoomed into show coverage of individual exons along with reads that span the intron indicated splicing.

## Future Directions

A major application of RNA-seq transcriptome is determining gene expression profiles. Currently, many such studies focus on identifying genes based upon the annotated genomic sequence for the target organism. However, RNA-seq data can provide a rich source of data for transcripts that are currently not recognized in a genome’s annotation file (Smith et al. current issue). Combining data from short-read (such as data generated by Illumina) and long-read (PacBio) RNA-seq technologies can provide an even better understanding of transcript structure, significantly improving genome annotation. Alternative splicing, allele specific expression, microRNAs and lncRNAs can all be identified with RNA-seq data, providing a rich catalog of the diversity of transcripts found in samples. Given the (relative) ease with which this data can be collected, an important current need is improving and extending bioinformatics tools. While many such tools exist, the community needs to work together to develop intuitive web-based bioinformatics pipelines and platforms that are appropriate to users with varying levels of computer sophistication. Ultimately, combining an incredibly rich set of sequence data with user-friendly bioinformatics tools will provide complete gene expression profiles of organisms in many different biological states.

## Acknowledgements

This material is based upon work supported by the National Science Foundation under Grant No. 1147029 and by the Agriculture and Food Research Initiative Competitive Grant No. by NIFA 2010-04233 from the USDA National Institute of Food and Agriculture.

